# Cryo-EM structure of the *Agrobacterium tumefaciens* type IV secretion system-associated T-pilus reveals stoichiometric protein-phospholipid assembly

**DOI:** 10.1101/2022.09.25.509369

**Authors:** Stefan Kreida, Akihiro Narita, Matthew D Johnson, Elitza I Tocheva, Anath Das, Debnath Ghosal, Grant J. Jensen

## Abstract

*Agrobacterium tumefaciens* is a plant pathogen that causes crown gall disease by the horizontal transfer of oncogenic DNA that is integrated into the host’s genome. The conjugation is mediated by the conjugative VirB/D4 type 4 secretion system (T4SS). *A. tumefaciens* T4SS assembles an extracellular filament, the T-pilus, that is involved in the formation of a mating pair between *A. tumefaciens* and the recipient plant cell by a not fully understood mechanism. Here, we present a 3 Å cryo-EM structure of the T-pilus, solved by helical reconstruction. Our structure reveals that the T-pilus comprises the major pilin protein VirB2 and phosphatidylglycerol (PG) phospholipid at a 1:1 stoichiometric ratio with 5-start helical symmetry. We further show that PG-headgroups and the positively charged Arg 91 residues of VirB2 protomers form extensive electrostatic interactions in the lumen of the T-pilus. Mutagenesis of Arg 91 destabilized the VirB2 protein and completely abolished pilus formation. While our T-pilus structure shows architectural similarity with previously published conjugative pili structures, positively charged sidechains protrude into the lumen and the lumen is narrower, raising questions whether the T-pilus is a conduit for ssDNA transfer. We also show that the VirB2 subunits in T-pilus filament are not cyclic, as previously thought.

## Introduction

*Agrobacterium tumefaciens* is a plant pathogen that, upon sensing plant wounds, horizontally transfers a segment of tumor-inducing (Ti-) plasmid DNA that is integrated into the host’s genome by nonhomologous recombination. Induction of these genes causes tumorous swelling of the plant tissue, known as crown gall, and the production of opines that can be taken up and catabolized by the bacterium (Van Attikum et al., 2001; Dessaux et al., 1998; Zhu et al., 2000). *A. tumefaciens* is routinely used in biotechnology to generate transgenic plants (Hwang et al., 2017), and more recently in the genetic manipulation of fungi (Idnurm et al., 2017). This plasmid transfer is an example of bacterial conjugation. Extensive study since its discovery in *Escherichia coli* (Lederberg and Tatum, 1946) has revealed that conjugation is widespread in prokaryotes and may be of narrow- (between the same bacterial species) or broad- (across phyla or kingdoms) host range (Virolle et al., 2020). Broadly speaking, conjugation allows prokaryotes to adapt to or, as in the case of the Ti-plasmid of *A. tumefaciens*, create a specific environmental niche, and thus conjugation plays an important role in bacterial evolution and pathogenicity (Norman et al., 2009).

Bacterial conjugation is mediated by the type IV secretion system (T4SS) (Costa et al., 2021). The conjugative T4SS in *A. tumefaciens* is encoded by the virulence (*vir*) region of the Ti plasmid and is comprised of 12 proteins (VirB1-VirB11 and VirD4) (Kuldau et al., 1990; Shirasu et al., 1990) that form an envelope-spanning complex and an extracellular filament (T-pilus). The T4SS is directly involved in mating-pair formation and transfer of Ti-plasmid DNA and effector proteins (Costa et al., 2021; Zhu et al., 2000). The T-pilus is primarily made up of VirB2 (major pilin) and decorated by a tip-protein, VirB5 (minor pilin) (Aly and Baron, 2007; Beijersbergen et al., 1994; Erh-Min and I., 1998). The exact role of the T-pilus in conjugation is unknown. However, a number of mutant alleles in VirB9-11 (reviewed in (Li and Christie, 2018)) have identified uncoupling phenotypes that are virulent (albeit reduced compared to wild type) but with no detectable pili (Vir^+^, Pil^-^). Functional VirB2 was required for virulence in the case of the uncoupling alleles in VirB9 and VirB11; and certain single-amino acid substitutions in VirB2 alone have resulted in the Vir^+^, Pil^-^ phenotype (Jakubowski et al., 2004, 2005, 2009; Wu et al., 2014). Together this suggests that VirB2 may have multiple roles (more than just being the T-pilus major pilin).

Previous structural and biochemical studies of three conjugative pili (pED208 of *Salmonella Typhi*, F-pilus of *E. coli* and pKpQIL of *Klebsiella pneumoniae*) have revealed that they are a 1:1 stoichiometric assemblies of protein and phosphatidylglycerol (PG) phospholipid. All three are so-called sex-pili that are involved in horizontal transfer of conjugative plasmids between bacterial species. They form hollow tube-like structures with outer diameters of 83-87 Å and luminal diameters of 25-28 Å, and with a net-negative luminal charge. The pED208 and F-pilus are highly homologous and are here referred to as F-like pili. The pKpQIL pilus differs mainly in rotational symmetry; *i.e.* the number of parallel helices in the filamentous structure. The F-like pili have 5-fold rotational symmetry while the pKpQIL is composed of one continuous pilus (1-fold symmetry) (Costa et al., 2016; Zheng et al., 2020).

The F-like pilin subunits are acetylated at the N-terminus following proteolytic cleavage of the propilin (Frost et al., 1983, 1984). T-pilins are reported to undergo a covalent cyclization between the N- and C-terminal residues of the mature pilin (Eisenbrandt et al., 1999). It is reasonable to assume that this difference may impact pilin packing in the assembled pilus. T-pili have been observed to be particularly resistant to harsh environmental conditions (Lai and Kado, 2002), but the structural basis remains unclear.

Here, using cryo-electron microscopy (cryo-EM), we reveal the structure of the T-pilus at 3.0 Å resolution. Our structure, consistent with other conjugative pili, reveals that the T-pilus is a 1:1 stoichiometric assembly of VirB2 pilin proteins and PG phospholipids. The structure is similar to the three reported structures in the near-luminal region that forms the hydrophobic core of the filament as well as the 5-fold symmetry that was shown for the F-like pili (Costa et al., 2016). However, our structure shows a neutral luminal charge with positively charged sidechains protruding towards the luminal core. Moreover, the lumen is narrower compared to other reported structures. Surprisingly, we do not observe cyclization of VirB2 subunits as expected.

## Results

### The T-pilus is a Five-start Helix

T-pilus filaments were obtained by growing the nopaline-type pTiC58-derived strain *A. tumefaciens* NT1REB(pJK270) (Chesnokova et al., 1997) in the presence of acetosyringone as described previously (Schmidt-Eisenlohr et al., 1999) (see also materials and methods). Purified filaments were analyzed by SDS-PAGE (Figure 1A). The gel band, corresponding to the size of processed pilin, 7.2 kDa (Shirasu and Kado, 1993) was excised and identified as major pilin VirB2 by in-gel digestion with chymotrypsin followed by mass-spectrometry with sequence coverage of the mature pilin.

**Figure 1.**
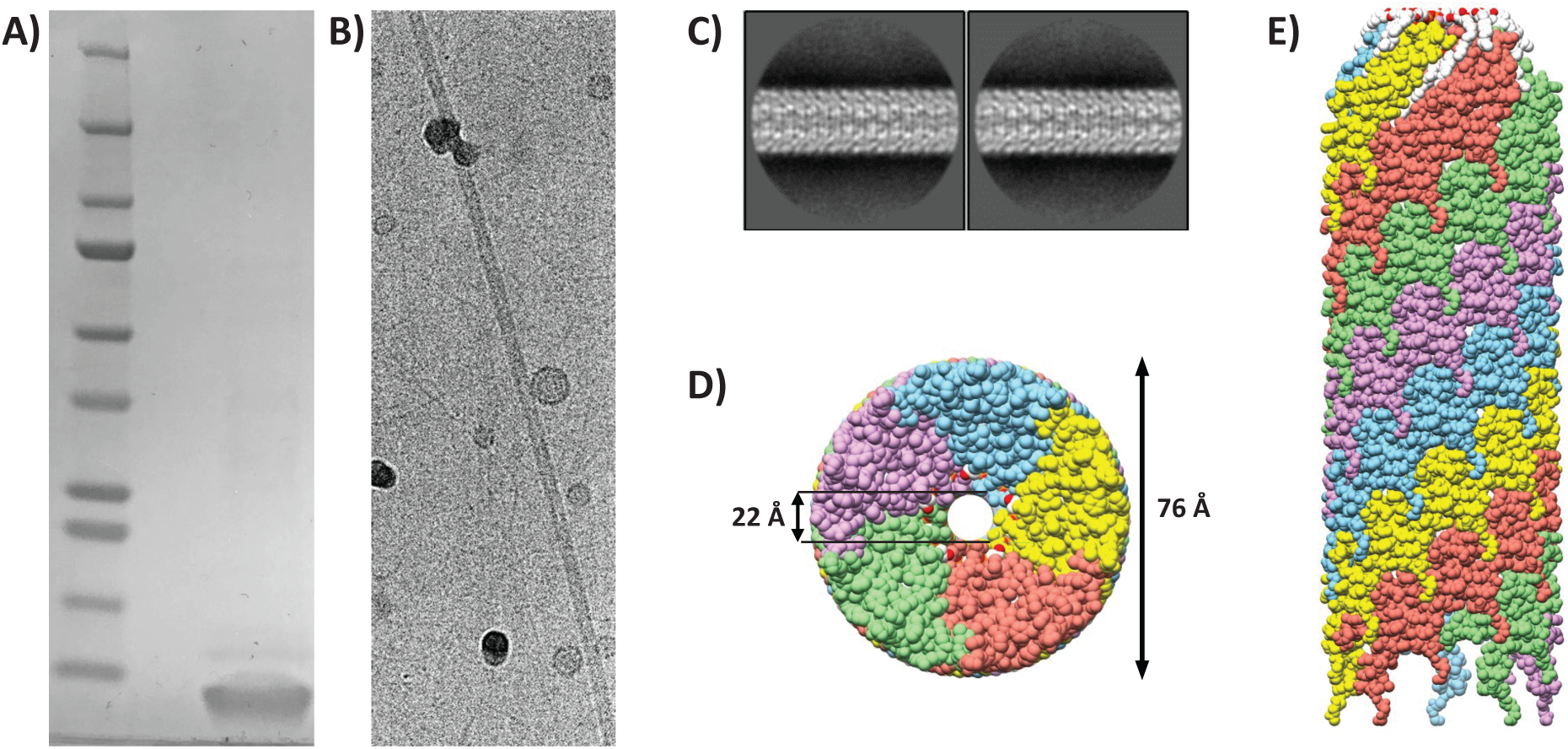
Structure determination of the T-pilus. **(A)** SDS-PAGE of purified VirB2 protein stained with Coomassie (lane 2). Mature VirB2 has a predicted MW of 7.2 kDa. Precision plus molecular weight marker was used (lane 1). **(B)** Cryo-electron micrograph of intact purified T-pilus **(C)** Representative 2D-class averages from Relion with a 300 pixel box size. **(D)** Bottom view of the filament model; coloured by individual VirB2 molecules. **(E)** Side-view of the filament. The filament is built up by five parallel helices, each of these have a unique colour. The top of the represented filament is towards the tip (as determined before (Meng et al., 2019)).

Cryo-EM data on purified pili were collected using a 300 keV Titan Krios microscope with a pixel size of 0.416 Å (super-resolution), later binned to 0.832 Å during processing. Electron micrographs revealed intact pili with lengths of >100 nm (Figure 1B). The data was processed within the Relion framework (He and Scheres, 2017; Scheres, 2012). Particles were extracted as overlapping boxes of 300 pixels with a shift of 14 Å and the structure was solved by helical reconstruction. The shift was an estimation of the helical rise by inspection of the 2D-class averages. Representative 2D class-averages are shown in Figure 1C. A total of 3543 boxes were used for the final reconstruction at 3.0 Å (map:map FSC at 0.143) (Figure S1 A-B, Table SI 1), with applied C5 symmetry (5 parallel helices), a rise of 13.4 Å and helical twist of 32.6°.

### The T-pilus is a tightly packed stoichiometric assembly of VirB2 and Phospholipid

The filament appears as a hollow tube with an outer diameter of 76 Å and luminal diameter of 22 Å (Figure 1D and E). The luminal diameter does not include Arg 91 sidechains, which protrude into the lumen but appear flexible. Segmentation of the map density showed that the filament is composed of two distinct densities that appear in the same stoichiometry (Figure 2A). Protein density dominated by alpha-helices fills most of the space and spans the filament at an angle (Figure 2B). This density was assigned to VirB2 (major pilin). The second density resembled a phospholipid (Figure 2A and C) with a small head-group appearing near the lumen and two hydrophobic tails extending along the VirB2 subunits. The lipid was identified as PG 16:1_18:1 (see Lipid Determination section below).

**Figure 2.**
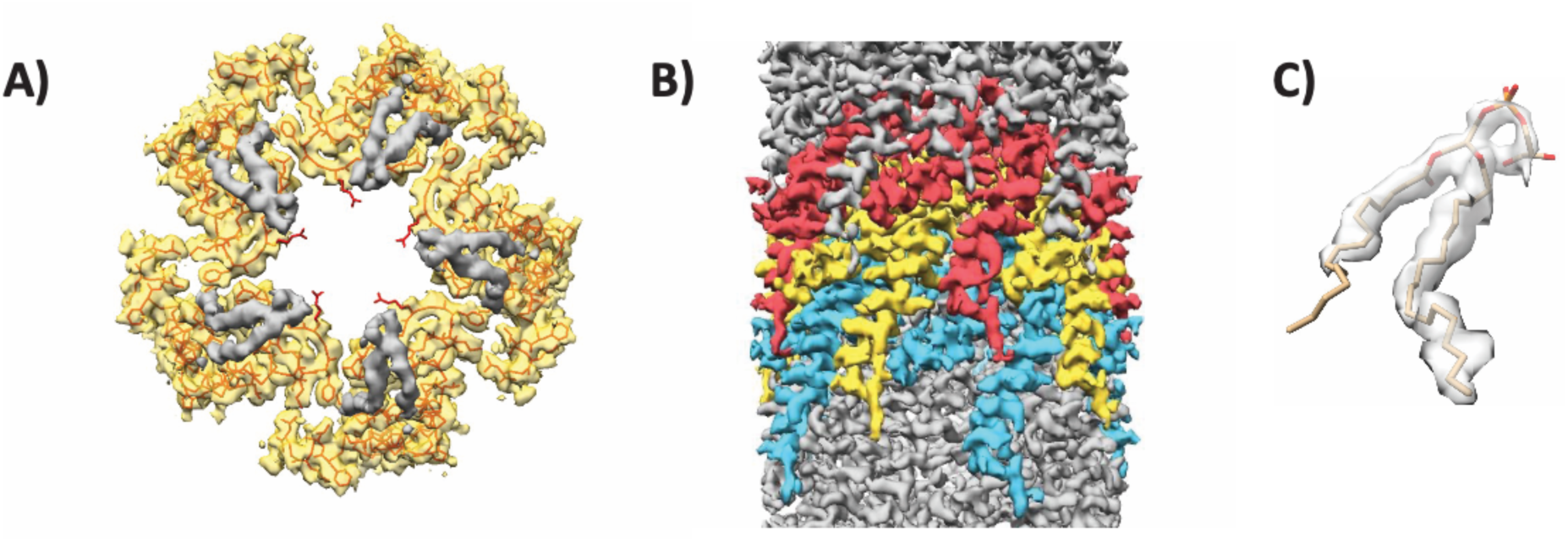
The T-pilus is composed of VirB2 and PG-phospholipid. **(A)** Map density of a helical layer as seen from the top. VirB2-assigned density in yellow with the protein chain represented as a string. Bulky sidechains are shown. PG-assigned density is coloured in grey. Sidechain of Arg 91 is seen extending into the lumen. **(B)** The filament seen from the side with three layers of VirB2. **(C)** Map/model coverage of a single PG molecule.

The T-pilus ultrastructure can be described as layered along the helical axis (Figure 3A and B). Each layer contains five copies of VirB2 and five phospholipids. As implied by the helical parameters, the layer is related to the layer above or below by a rotational angle of 32.6° and a rise of 13.4 Å (Figure 3C-D). The rotation positions the phospholipids of layer n+1 (towards the pilus tip) directly above VirB2 protomers of layer n. The result of these helical operations is that each VirB2 subunit has a large interaction surface with acyl chains of four surrounding phospholipid molecules (Figure 4A-C). Similarly, each phospholipid molecule interacts with four neighboring VirB2 monomers forming an intricate interaction network. Moreover, each VirB2 protomer forms extensive hydrophobic interactions, as well as hydrogen bonds via Asn 62 and Gln 73 to neighboring VirB2 protomers (Figure 4D).

**Figure 3.**
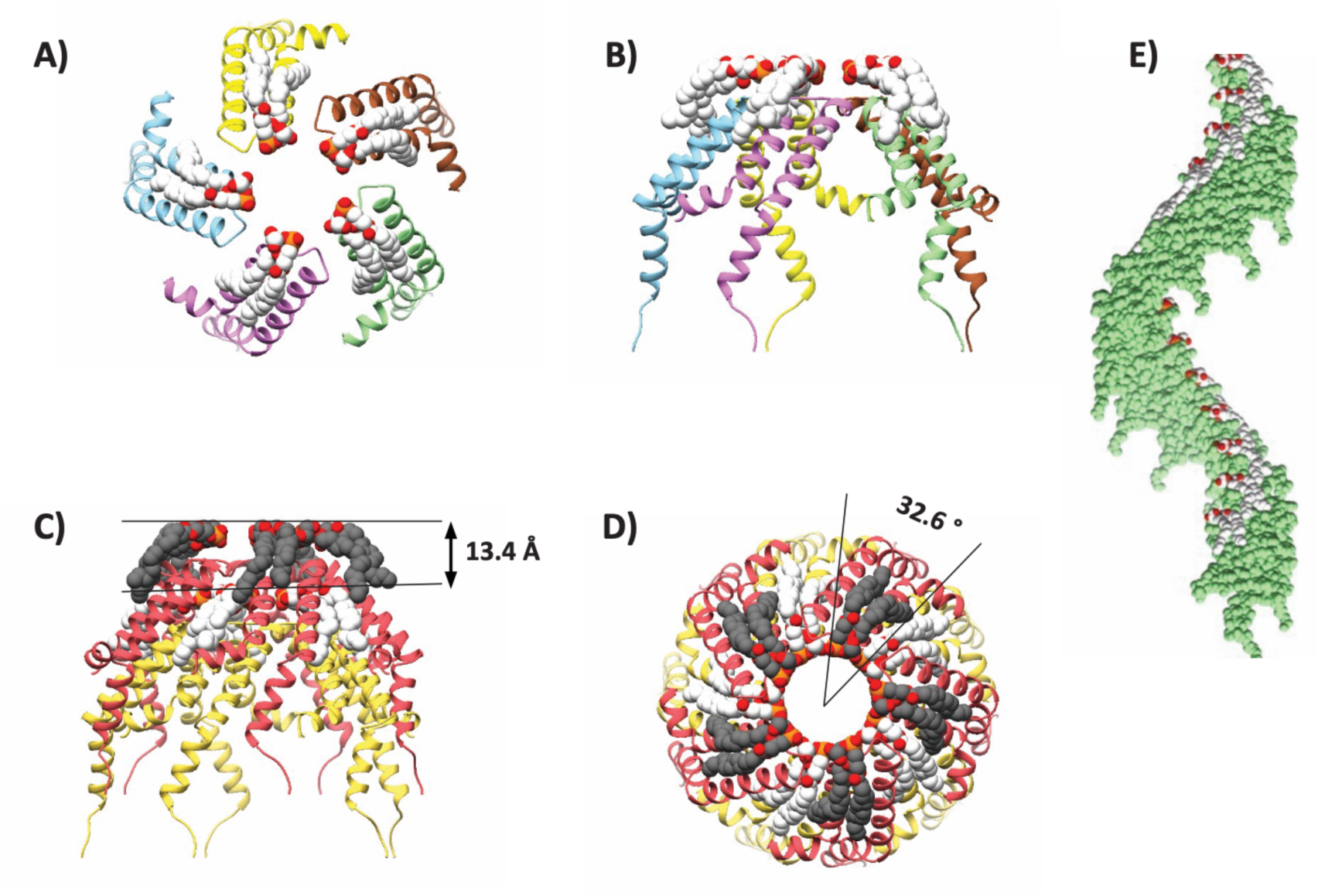
Helical symmetry of T-pilus. **(A-B)** Ribbon representation of the top and side view of a helical layer. **(C-D)** Representation of the helical rise and twist. **(E)** Helical axis of a single helix in the symmetry.

**Figure 4.**
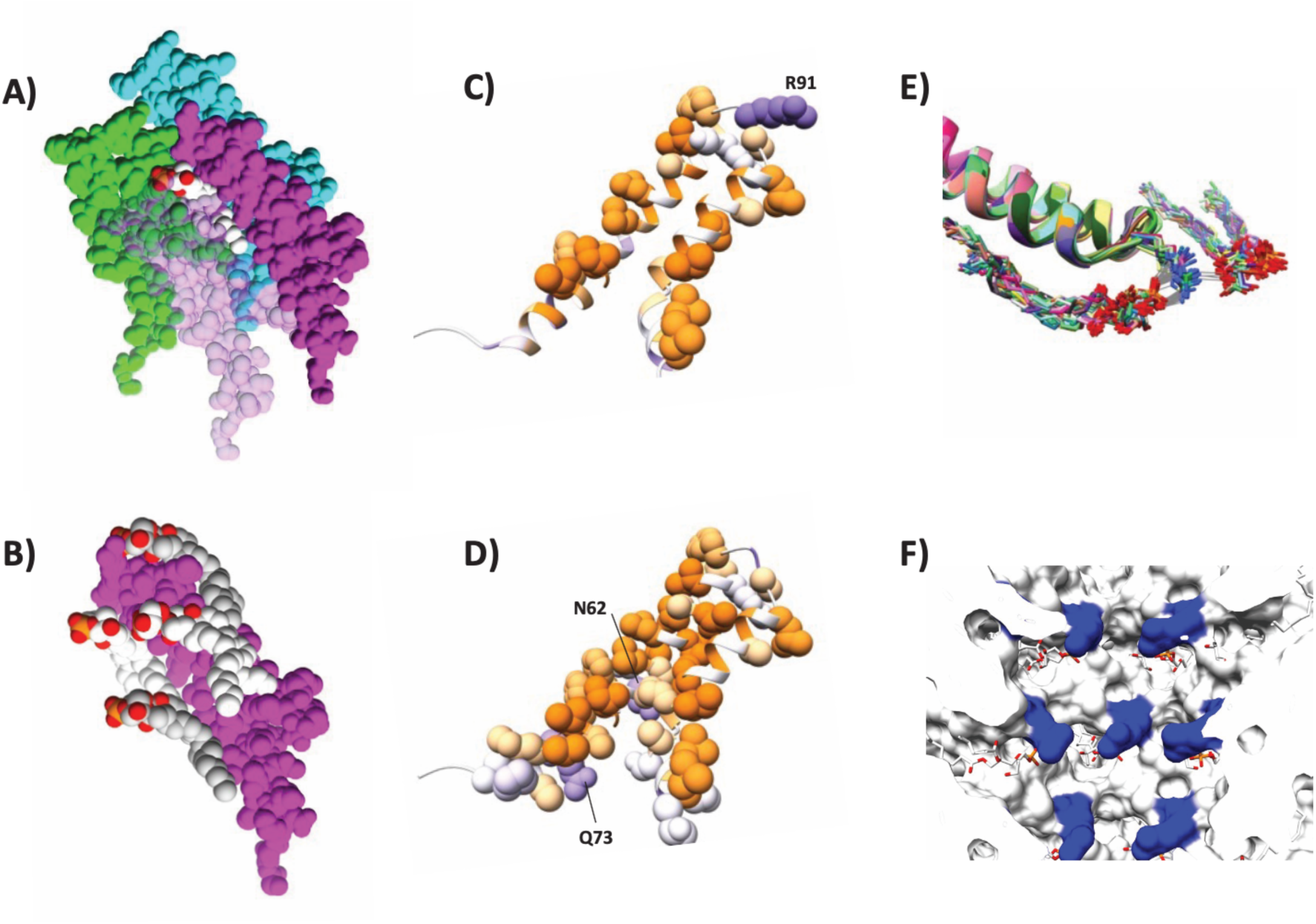
Interaction network between virB2 monomers and lipid molecules. Space filling models showing that **(A)** each PG molecule (white) interacts with four VirB2 protomers (purple, cyan, green and light pink) and **(B)** each VirB2 protomer (purple interacts with four PG molecules (white and light-grey). VirB2 Interface involved in **(C)** protein-lipid or **(D)** protein-protein interaction, colour-scaled by hydrophobicity (purple: hydrophilic, orange: hydrophobic). **(E)** Flexibility of Arg 91 as determined by MD simulations showing that it is possible for Arg 91 sidechains to interact with two lipidic head groups by electrostatic interaction and hydrogen bonds. 30 unfixed molecules from two independent MD simulations are overlayed. Grey lines indicate hydrogen bonds between lipids and VirB2 protomers. **(F)** View of the pilus lumen showing Arg 91 (blue) and phospholipids (white/red).

The VirB2 chains are somewhat U-shaped. The termini are located on the outward side of the filament and are not covalently joined – *i.e.* the VirB2 subunits are not cyclic in our structure as previously reported from mass spectrometry (Eisenbrandt et al., 1999). Starting from the N-terminus, the chain runs towards the lumen and forms two helices - α1 and α2 (Figure S2 A). The chain turns in the lumen and returns outwards via two helices α3a and α3b. As shown in Figure 4, residues along the length of each protein chain are involved in inter-protein or inter-lipid interactions with multiple proteins and lipids. α2 and α3a are anti-parallel to each other, largely hydrophobic and buried in the structure. The residues separating the two helices form a luminal loop (residues 89-93) that holds Arg 91, the only positively charged residue in the lumen (Figure 4C,E-F). The position of the Arg 91 sidechain is not well resolved the map density (Figure 2A). Molecular dynamics simulations (see Materials and Methods) showed that the position of the arginine sidechain fluctuates so that the guanidino groups can interact with two separate phosphate groups (the only negatively charged functional groups in the lumen) in the helical assembly (Figure 4E). This results in a network of electrostatic interactions in the lumen with no net charge (Figure 4F, Figure S2 B-C).

### The Luminal Loop is important for T-pilus formation and Virulence

To investigate the importance of the VirB2 luminal loop, we generated mutant VirB2 proteins containing individual R91E, R91A and S93A mutations. We tested these mutants for their ability to form an active pilus using western blot and functional assays (Figure 5A). Mutant versions of VirB2 were expressed in *A. tumefaciens* PC1002 that contains a nonpolar deletion in *virB2*. Expression was monitored by Western blot using anti-VirB2 specific antibodies. The western blot showed no detectable band for R91E and R91A, indicating that these mutations destabilized VirB2 (Figure 5A). VirB2 S93A mutant was stable but accumulated at a lower level. Analysis with anti-VirB10 antibody showed that *vir* genes were induced in all mutants. Negative staining of cells grown under *vir*-inducing conditions showed that wild type cells displayed T-pili, whereas no pili were observed in R91A (n=100). Representative examples are shown in Figure S3.

**Figure 5.**
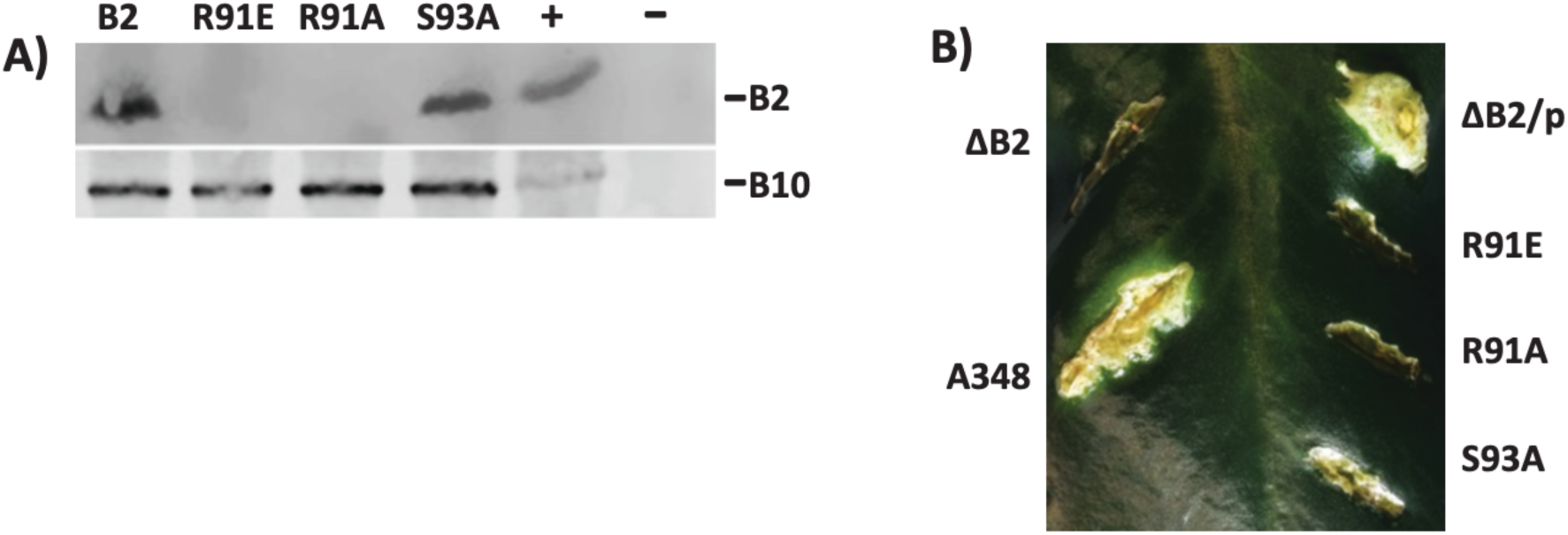
Importance of luminal loop in structure and function of the T-pilus. **(A)** Analysis of VirB2 mutant protein accumulation by western blot assays. *A. tumefaciens* expressing *virB2* or its mutants were analyzed by SDS-PAGE followed by western blot assays using anti-VirB2 antibodies. The blot was reprobed with anti-VirB10 antibodies as a control for induction of the *vir* genes. -, -Ti plasmid; +, pTi. **(B)** Effect on mutations in VirB2 Arginine 91 and Serine 93 on *A. tumefaciens* virulence. The effect of a mutation in *virB2* was monitored by complementation assays. *Agrobacterium* PC1002 (ΔB2) and its derivatives harbouring a plasmid that expressed *virB2* (p) or its mutant derivatives B2R91E (R91E), B2R91A (R91A) or B2S93A (S93A) were used to infect *Kalanchoe daigremo*ntiana leaves.

The *Kalanchoe daigremontiana* leaf infection assay is a standard functional assay used for monitoring Vir protein function. The results of the functional assays correlated well with the pilus protein expression levels. Strains with VirB2 mutants R91E and R91A failed to form tumors indicating both mutants were non-functional in DNA transfer. The VirB2 S93A mutant is functional since bacteria producing this protein showed reduced virulence (Figure 5B). The reduced virulence can be due to its lower abundance in the cell and/or a negative effect on its function.

### The T-pilus incorporates PG

The lipid was identified by shotgun lipidomics spectrometry, in which the lipid content of purified pili was compared to that of a negative control. The negative control was produced by growing *A. tumefaciens* under non-inducing conditions and subjecting the cells to the same purification procedure as for the pili-containing sample. Samples were analyzed in triplicates from three independent preparations and the presence or absence of pili was verified by negative staining.

Lipids were extracted from the sample using a modified Bligh and Dyer extraction procedure and were analyzed for lipid content (Hsieh et al., 2021; Su et al., 2021). Lipid standards of 1450 lipid species were added to annotate lipids in biological samples. An independent experiment in which different concentrations of the same preparation (pili and negative control, respectively) were analyzed for lipid content (data not shown). Lipid species that did not increase in abundance with increasing sample concentration were considered contaminants (*e.g.* from tubes, lipid extraction solutions) and disregarded from analysis. The analysis yielded 24 non-contaminant lipid species in the samples, all belonging to PE, PG and PC classes. (Table SI 2).

If the T-pilus itself incorporates a lipid, it would be expected that a specific lipid species would increase as a fraction of the total composition in the pili-containing sample compared to the negative control. PG 16:1_18:1 stands out from the 24 selected lipid species as it shows a 9.2-fold increase in the pili-containing sample (MANOVA, p<0.001) (Figure S4A). It was necessary to mathematically quantify how important this increase was compared to all changes in the composition to a draw a conclusion. Unaveraged data was therefore transformed to centered log-ratios (CLRs) and the variance was calculated for each lipid species from all 6 observations (3 from pili and 3 from negative control). A large difference between induced and non-induced cells would thus result in a high variance. The variance of each lipid species was then divided by the sum of that of all species and normalized to 100% (contribution to total variance) (Figure S4B). This metric showed that the observed increase in PG 16:1_18:1 accounted for 37% of the total variance in the dataset, which is large considering that 24 lipids species were included. The analysis further revealed that PG 18:1_18:1 increases 3-fold (p<0.001) and that this increase accounted for 13% of the total variance in the dataset. Since PG 16:1_18:1 and PG 18:1_18:1 are chemically similar, a model in which the T-pilus incorporates both species is likely and would explain 50% of the total variance in the dataset. We therefore propose that the T-pilus is promiscuous and incorporates both these lipids, but preferentially PG 16:1_18:1, which was modelled into the structure.

## Discussion

Conjugative pili have been classified by morphology and general functional characteristics. So-called F-type pili (e.g., pED208, F-pilus) and pKpQIL-pili are characterized by thin, long and flexible filaments that promote efficient conjugation in both solid and liquid environments and have a narrow host-range. In contrast, the P-type pili (which includes the T-pilus) are generally thicker, shorter and more rigid, are more efficient in solid environments and have a broader host-range (Christie, 2016; María and Fernando, 2018). These traits should be seen as general trends that reveal evolutionary origins, but exceptions exist, *e.g.,* the T-pilus is thinner than the F-pilus and forms long filaments (Figure S3).

Our near-atomic resolution structure of the T-pilus revealed that it is a stoichiometric assembly of protein and PG phospholipid, similar to the F-type pili (pED208, F-pilus and pKpQIL conjugative plasmid-derived pili), described earlier (Costa et al., 2016; Zheng et al., 2020). Since this is the fourth conjugative pilus structure solved to date, it seems likely that all conjugative T4SS pili are stoichiometric protein-lipid assemblies. The phospholipids have a clear structural role in the T-pilus and can be likened to putty that fills the gaps between VirB2 subunits. Moreover, positively charged residues in the luminal loop (a conserved feature in VirB2/TraA pilins (Lai and Kado, 2000)) and the negatively charged phospholipid headgroups create a network of alternating positive and negative charge that may have a function in stabilizing the structure of the lumen. Beside this structural role, we speculate that the lipids may play functional roles in establishing contact (or fusing) with the target membrane or facilitate movement of pilin subunits back into the membrane during pilus retraction. This should be the subject of future investigations.

VirB2 adopts a distinct fold with structural homology to that of the other known conjugative pili structures. Structural comparison between the T- and F-pili (PDB ID: 5LEG) revealed that they are architecturally similar in the near-luminal and buried regions composed of the parallel helices α2 and α3 (Figure 6). However, VirB2 has an elongated α1 and a 72° kink in α3 at Gly 109. G109A substitution was previously shown to abolish pilus formation and virulence (Wu et al., 2014), suggesting that division of α3 into two helices (here called α3a and α3b) is important for T-pilus stability. The structural difference between T- and F-pili pilins likely causes the difference in packing of the assembled filament. The result is that the T-pilus is denser (with smaller filament and luminal diameters) than the F-pilus despite the subunits being of similar size and molecular weight. The luminal diameters of the F-family pili are 28 Å, whereas that of the T-pilus was 22 Å, not including the Arg 91 sidechains. The net charge of the lumen is also different. Whereas known conjugative pili have a net-negative luminal charge, the T-pilus has a net neutral charge, but with positively charged sidechains extending towards the center of the lumen (Figure S2). As the positive charge would attract the negatively charged ssDNA, it is unclear whether this would facilitate ssDNA transfer.

**Figure 6.**
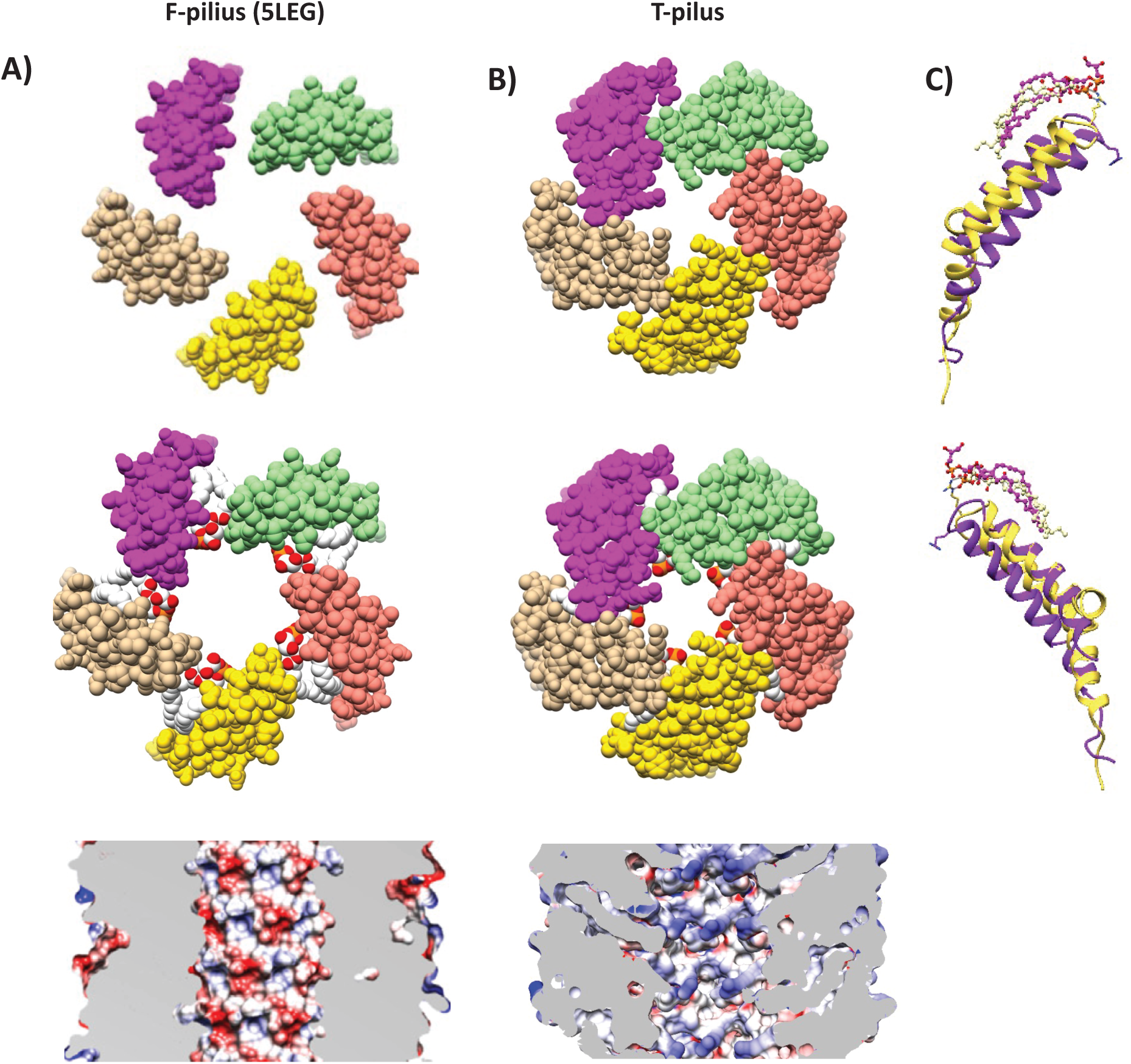
Structural Comparison of the T- and F-pilus. Helical cross-sections showing organization of pilins and phospholipids in (A) F-pilus (PDB ID: 5LEG) and (B) T-pilus. The T-pilus adopts a denser structure with narrower lumen. The F-pilus lumen is moderately negatively charged (reproduced from Costa et al., 2016), whereas T-pilus lumen is neutral. (C) Structural overlay and secondary structure annotation of T-pilus VirB2 (yellow) and F-pilus TraA (purple). Phospholipids from equivalent positions in the helical packing are displayed.

Surprisingly, VirB2 did not form cyclic subunits as seen previously by mass-spec of purified T-pili (Eisenbrandt et al., 1999). The first and last three residues of mature VirB2 were not modeled due to poor map density at the termini, but the modelled termini were ∼30 Å apart in the structure - too far to be connected by a cyclic linkage. Cyclic VirB2 may thus represent an intermediate form that is later linearized or, as VirB2 appears to have multiple roles during mating pair formation, a different functional form.

The T-pilus has 5-fold rotational symmetry, just like pED208 and F-pilus, but different from the pKpQIL conjugative pilus. The recent cryo-EM structure of the R388 conjugative T4SS (Macé et al., 2022) exhibited a previously unseen stalk complex in the periplasm between the inner- and outer membrane core complexes (OMCC). The stalk was made up of pentamers of VirB6 and thus matched the 5-fold rotational symmetry of the F-like pili. A model was therefore proposed that the pilus is formed on top of the VirB6 platform, with the pentameric VirB5 tip being pushed up through the OMCC by polymerization of VirB2. This dictates the filament must be narrower than the periplasmic cavity of the OMCC pore to be able to fit and be 5-fold symmetric. The OMCC of VirB/D4 was previously determined to have an inner diameter of 82 Å (Gordon et al., 2017). The T-pilus has the required 5-fold symmetry and diameter of only 76 Å. The structure is thus in agreement with the model of pilus assembly proposed by Macé et al. (2022).

The role of the luminal Arg 91 is interesting. The model presented here suggests that it has a structural role in creating a patchwork of electrostatic interactions with the lipid headgroups. It has previously been shown that R91A substitution results in loss of virulence and deficiency of VirB2 pre-pilins, suggesting that it has a role in folding in the inner membrane (Wu et al., 2014). These findings are supported by our *K. daigremontiana* leaf infection assay. Substitution of Arg 91 to either alanine or glutamic acid resulted in no detectable pili or virulence. Furthermore, S93A substitution, a residue of the luminal loop that is not involved in lipid interaction, resulted in reduced virulence and reduced extracellular pilus formation. These effects could be attributed to structural changes of the luminal loop, which effect helical packing (as was observed in (Zheng et al., 2020) and T-pilus stability.

PG was also found in the F-family pili, suggesting it to be a conserved feature in conjugative T4SS pili. In support of this identification, *A. tumefaciens* contains two virulence genes, *lpiA* (a Lysyl-PG synthase) and *acvB/VirJ* (Lysyl-PG hydrolases), that are involved in PG maturation in the inner membrane (Groenewold et al., 2019; Pan et al., 1995; Wirawan et al., 1993). In C58-derived *A. tumefaciens* strains these genes are located chromosomally in the same operon and are, like the T-pilus machine, both induced at low pH (McCullen and Binns, 2006; Vinuesa et al., 2003; Winans, 1990). Moreover, the octopoine pTiA6 plasmid contains *VirJ* (an *acvB* homologue) directly under the control of VirA/VirG, the two-component system that is induced by certain plant phenols (e.g acetosyringotone) and controls the transcription of *Vir* genes (Kalogeraki and Winans, 1995). These PG-maturation genes are thus co-induced with the T-pilus machine. Interestingly, *acvB* inactivation was shown not to affect cellular levels of VirB8 and VirB9 (used as markers for T4SS) but reduced levels of the minor pilin VirB5 (VirB2 levels were not monitored). These findings together suggest that PG maturation has an impact on pilus formation, which is explained by our findings. We note that VirA/VirG-induced PG maturation and VirB2 folding both occur in the inner membrane and VirB2 extraction was proposed to be mediated by VirB4 (Groenewold et al., 2019; Kerr and Christie, 2010). It is thus possible that VirB2 and PG are removed from the inner membrane together for assembly into the pilus in a 1:1 ratio. Phosphatidylcholine (PC) synthase has also been identified as a critical virulence factor (Aktas et al., 2014). However, PC-deficient mutants did not produce T4SS machines and showed no sensitivity to acetosyringone (Klüsener et al., 2010), suggesting that PC-deficiency has an upstream effect on VirA/VirG regulation of the *vir* operon.

In a contemporary study, Amro *et al* also resolved the structure of the *A. tumefaciens* T-pilus at comparable 3.2 Å resolution (Amro et al., 2022). While their structure and ours are very similar, they identified the lipid as PC. PC species were present in the pili-containing fraction of our experiment, but their part of the composition did not increase substantially compared to the negative control (Table S2), causing us to reject PC as a candidate for the lipid. Notably, PG species were not identified at all in *A. tumefaciens* membranes in this study; however, *A. tumefaciens* has been shown to contain PG species (Aktas et al., 2014 and this study), and as stated above, holds genes for PG maturation. The absence of PC in their study may therefore be due to strain-related differences, or perhaps their experimental setup did not allow detection of PG.

Pili make part of prokaryotes’ toolbox to interact with their environment. While the similarities of the T-pilus structure presented here with that of other conjugative pili is striking and suggest a conserved function, the differences may indicate interesting specializations.

## Materials and Methods

### A. tumefaciens Growth

*A. tumefaciens* cells were grown and purified under conditions similar to those described previously (Schmidt-Eisenlohr et al., 1999). An overnight culture in LB was used to inoculate AB/pH 5.5 buffer (1 gL^-1^ NH_4_Cl, 0.3 gL^-1^ MgSO_4_ 7 x H_2_O, 0.15 gL^-1^ KCl, 10 mgL^-1^ CaCl_2_, 2.5 mgL^-1^ FeSO_4_ 7xH2O, 3.9 gL^-1^ MES, 10 gL^-1^ Glucose, 10 mM Na/KPO_4_, pH 5.5) to an OD^600^ of 0.1. Cells were grown at 28°C in a shake-incubator at 220 rpm. For (non-induced) negative control, 2.4 L of culture was grown for 72 h. For induced cells, a 100 mL of culture was grown for 5 h, then centrifuged at 4,000 x g (20 min) at room temperature and the pellet resuspended in ¼ of the original volume in fresh AB/pH 5.5 buffer. 0.5 mL of this suspension was spread per each 15 mm petri dishes containing AB/pH 5.5 buffer, 1.5% Bacto Agar (Difco), 300 μM acetosyringone (Sigma). 40 such plates were made for each preparation. The plates were incubated at 22°C (72 hours).

### Pili Purification for Cryo-EM and Lipid Analysis

The nopaline-type pTiC58-derived strain *A. tumefaciens* NT1REB(pJK270) (Chesnokova et al., 1997) was used for pili purification. Cells were grown as described above. Induced and non-induced cells were harvested by centrifugation at 6,000 x g at 4°C. Induced cells were scraped from plates and resuspended in 50 mM MES, pH 5.5 prior to centrifugation.

Cells were resuspended in 12 mL 50 mM MES and passed through a 26-gauge syringe needle ten times in and out. This procedure causes the cells to shed its pili. The suspension was centrifuged at 10,000 x g at 4°C (1 h). The supernatant was then ultracentrifuged at 100,000 x g (1.5 h). The resulting pellet was resuspended in 50 mM MES pH 5.5. The pilin was identified as VirB2 by mass-spec (Q-Exactive HF, LCMS) at the Caltech Proteome Exploration Laboratory. Presence or absence of pilus filaments was confirmed by negative staining on formvar grids with 3% uranyl acetate (30 s).

### Sample Preparation and Data Collection for Cryo-EM

Pili sample was cryogenically frozen on a FEI Vitrobot MK4, set to 22 °C and 100% humidity. 3 uL of sample was blotted on glow-discharged UltraAuFoil EM-grids (R1.2/1.3, 300 mesh, Electron Microscopy Sciences), with 3 s of wait time and 7 sec of blot time. The data was collected at the Caltech Cryo-EM facility on a 300 kV FEI Titan Krios equipped with a Gatan K3 direct detector and Gatan energy filter. Movies of 40 frames were collected at a calibrated pixel size of 0.416 Å (super res), with a total dose of 60 e^-^/Å^2^. The defocus range was set to -0.6 to -2.0.

### Helical Reconstruction and Data Processing

The data was processed in Relion by helical reconstruction (He and Scheres, 2017; Scheres, 2012), using Motioncorr2 for motion correction (Zheng et al., 2017) and CTFFIND-4.1 (Rohou and Grigorieff, 2015) for CTF-estimation. The data was downsized to 0.832 Å and overlapping 300×300 pixel boxes were extracted with a 14 Å shift (∼equivalent to a helical rise). A 7 nm cylinder was use as a starting model for 3D classification. The initial helical rise and pitch were estimated using the 2D class averages. 3D classification was then used to screen the different combination of twist and n-fold symmetry that would be possible given the estimated helical rise and pitch. The highest resolving 3D class averages were 3D-refined and visually inspected in Chimera for features expected at the given resolution. 3543 boxes were used in the final reconstruction with a rise of 13.4 Å, helical twist of 32.6° and C5 symmetry.

### Model Building and Molecular Dynamics Simulations

The Initial model was generated with SWISS-MODEL (Guex et al., 2009) from the mature VirB2 sequence and fitted to the density map using Rosetta3 (Leaver-Fay et al., 2011). The resulting model was iteratively refined with COOT (Emsley et al., 2010); molecular dynamics flexible fitting (MDFF), using the ChimeraX extension ISOLDE (Croll, 2018; Goddard et al., 2018); and Phenix (Adams et al., 2011).

Molecular Dynamics simulations were performed in NAMD (Phillips et al., 2020) to estimate the dynamics of the Arg 91 sidechain. The simulations included 5 layers of proteins and lipids, explicit waters and 100 mM NaCl. The top and bottom layers were fixed. All C_α_ atoms in the protein, except for the five residues surrounding Arg 91; and all non-H atoms in the lipids were under weak harmonic constraints with k = 0.3 kcal mol^-1^ Å^-2^. The simulations were performed at 300 K for 1 nsec. Two independent simulations were performed.

### *virB2* Mutagenesis

Mutations in *virB2* were introduced using QuikChange II site-directed mutagenesis kit (Agilent Technologies). Plasmid pAD1891 DNA, a pUC118 derivative with a *virD*p-*virB2* chimeric gene was used as template DNA. Mutations were introduced by PCR amplification using primer pairs (mutations in bold):

GTTCGGG**GA**GGCTTCGCTTGGGCTG and GCGAAGCC**TC**CCCGAACATCCAGGAG (R91E); GTTCGGG**GC**GGCTTCGCTTGGGC and GCGAAGCC**GC**CCCGAACATCCAGGAG (R91A) or GGGCGGGC**AG**CGCTTGGGCTGGTTG and CCCAAGCG**CT**GCCCGCCCGAACATCC (S93A).

The mutations were confirmed by DNA sequence analysis of the entire genes. The plasmids were fused to the plasmid pAD1412 (Mossey et al., 2010) to construct the wide-host-range derivatives. Plasmids were introduced into *Agrobacterium* PC1002 by electroporation. PC1002, a derivative of the wild-type *Agrobacterium* strain A348 that carries the octopine type Ti-plasmid pTiA6NC, has a nonpolar deletion in *virB2* (Berger and Christie, 1994).

### Protein analysis and functional assays

Complementation assays were used to monitor the effect of a mutation on *virB2* function (Mossey et al., 2010). *Agrobacterium* PC1002 derivatives harboring a plasmid expressing *virB2* or its mutant were used to infect *Kalanchoe daigremontiana* leaves. Tumor formation was monitored 3 weeks after infection.

Accumulation of VirB2 mutant proteins was analyzed by western blot assays (Mossey et al., 2010). Total proteins from induced *Agrobacterium* strains were separated by SDS-polyacrylamide gel electrophoresis, transferred to a nitrocellulose membrane and probed with anti-VirB2 antibodies.

### Negative Staining and Imaging of A. tumefaciens Cells

*A. tumefaciens* cells were grown on plates in presence of acetosyringone as described above. Cells were collected after 72 hours using a sterile cotton swab and resuspended in 50 mM MES pH 5.5. Cells were not centrifuged to prevent pili breaking from the cells. 3 μL of this suspension was deposited on glow-discharged 300 mesh formvar/carbon copper grids (Electron Microscopy Sciences), washed once with water and stained with 0.5% w/v phosphotungstic acid, pH 5.5, for 30 s. Cells were imaged on a 120 kV FEI Tecnei Ti12 with LAB6 filament and Gatan Ultrascan 2k x 2k CCD.

### Lipid Data Collection, selection and treatment

#### Lipid Extraction and Mass Spectrometry Lipidomics Analysis

Lipid Extraction and mass spectrometry was performed by UCLA Lipidomics. A modified Bligh and Dyer extraction (Hsieh et al., 2021) was done to extract lipids from samples. Prior to extraction, 70 lipid standards across 17 subclasses were added to each sample (AB Sciex 5040156, Avanti 330827, Avanti 330830, Avanti, 791642). Extraction was performed twice, and the pooled organic layers were dried by Speedvac (Thermo, SPD300DDA) with ramp setting 4, 35°C for 45 min with a total run time of 90 min. Samples were resuspended in 1:1 methanol/dichloromethane with 10 mM ammonium acetate and transferred to robovials (Thermo, 10800107) for analysis.

Samples were analyzed by direct infusion on a Sciex 5500 with differential mobility device (DMS), tuned with EquiSPLASH LIPIDOMIX (Avanti 330731) with a targeted acquisition list of 1450 lipid species. Descriptions of instruments settings, MRM list and in-house data analysis workflow is available (Su et al., 2021).

### Lipid selection and Lipid Data Analysis

Three independently purified samples of purified pili and negative control were analyzed for lipid content. Lipids were selected for subsequent analysis by an independent experiment, in which dilution series (1x, 0.5x, 0.25x) of induced and non-induced pili preparation were analyzed for lipid content. Lipid species were selected if they increased in abundance (in nmoles) with increasing sample concentration. Some lipid species were not annotated in the selection experiment but appeared in different amounts in induced and non-induced cells and were included in the analysis. These were: PE 14:0_16:3, 14:0_18.2, 14:1_18:2, 15:0_18.2, 15:0_20.5, 16:0_16:2, 16:0_16:3, 16:0_17.1. The total number of selected lipid species were 24.

4 out of 144 measurements (2.78%) were missing values and needed to be generated to be able to log-transform the data for subsequent multivariate statistical analysis. The missing values were in species PE 14:0_16:2, PE 14:0_16:3 and PE 16:0_17:1. Visual inspection of the raw data showed that the main variation was between, rather than within triplicates of induced and non-induced cells. Moreover, existing data of affected lipid species did not suggest that the missing values were missing because of being below the detection limit. Replacing the missing values with low positive values would thus risk introducing false positives. Instead, they were estimated using existing measurements within the triplicates using the R-package zCompositions (Palarea-Albaladejo and Martín-Fernández, 2015). SD was not calculated for these lipid species.

Data was treated as compositional and closed to 100% (table 1). For MANOVA and variance calculations, data was first transformed to centered logratios (CLRs) (Greenacre, 2021). with weights proportional to species abundance (giving higher weight to more abundant species in the compositions). Variance was calculated for each lipid species as the variance of all observations (induced and non-induced cells, n = 6), divided by the sum of variance of all species and closed to 100%. MANOVA was performed in R (R Core Team, 2013). CLR transformation and variance calculations were done using the R package easyCODA (Greenacre, 2021). Plots were produced in ggplot2 (Wickham, 2016).

## Acknowledgements

We thank Dr. Songye Chen, Caltech cryo-EM facility for assistance during data collection; Dr. Tsui-Fen Chou, Dr. Brett Lomenick and Dr. Jeff Jones from Caltech Proteome Exploration Laboratory for conducting the protein mass-spec analysis. We also thank Dr. Kevin Williams and UCLA Lipidomics for performing lipid extraction, lipid data collection and giving valuable advice in lipidomic experimental design and data interpretation. This project was funded by a National Institutes of Health grant (R01 AI127401 to G.J.J), a National Health and Medical Research Council grant (APP1196924 to DG) and a Natural Sciences and Engineering Research Council of Canada Discovery grant (RGPIN 04345 to EIT). SK is supported by the Swedish Research Council (2019-06293)

## Supplementary Information

**Figure SI 1.**
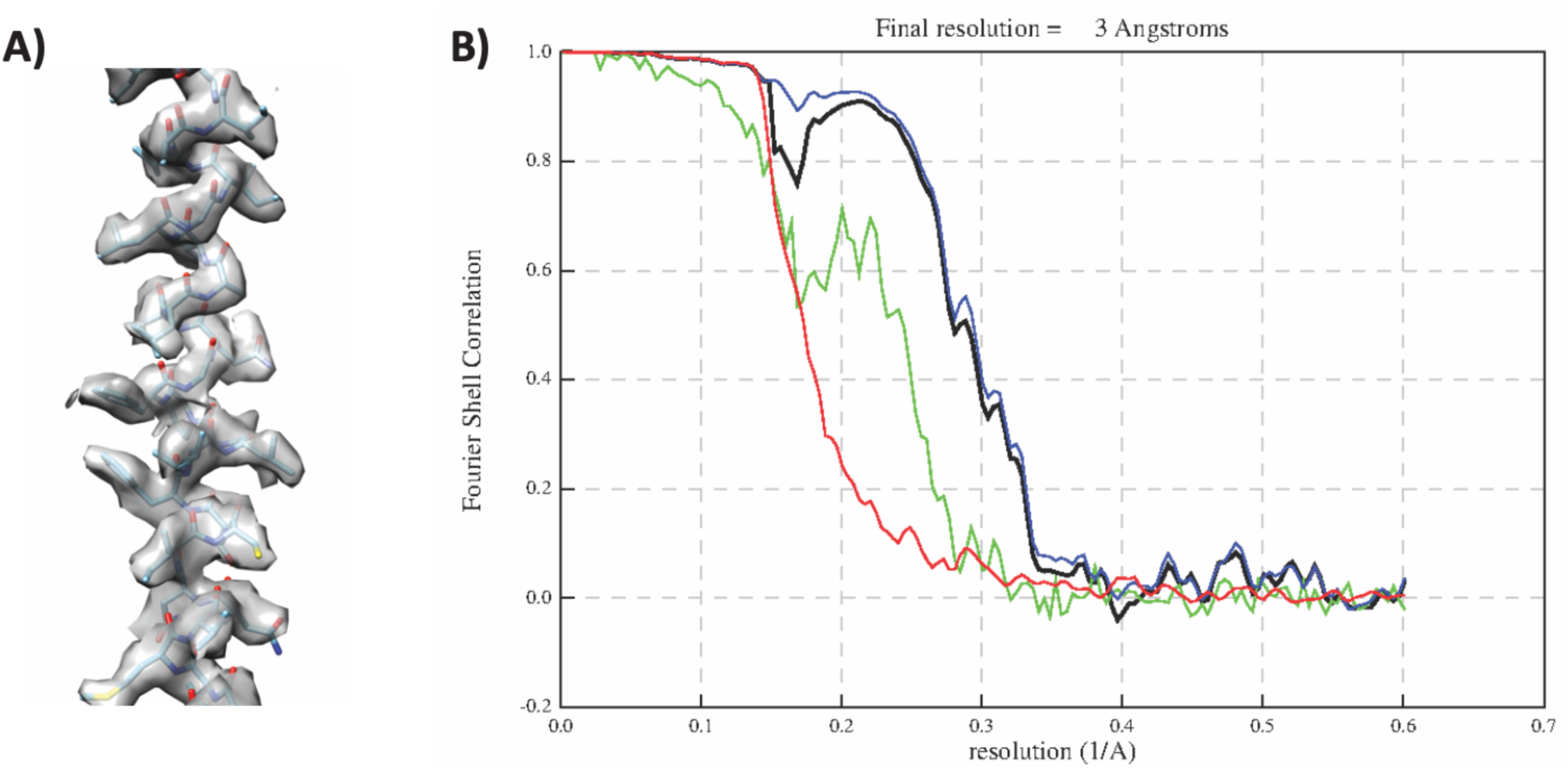
Resolution estimation. (A) Representative region of VirB2 monomer showing the fit of the atomic model to the experimentally derived cryo-EM density map. (B) Fourier shell correlations (FSCs) for evaluating the resolution calculated by RELION (Scheres, 2012). FSCs between two 3D structures from each half of the dataset with or without masking are presented in green and blue, respectively. The red curve was calculated after the phase was randomized beyond 6.9 Å to evaluate the artifacts from overfitting. The black curve is the corrected FSC after accounting for the artifacts from overfitting. Based on the gold-standard criteria between two half-maps, the resolution is 3.0 Å with a threshold of 0.143.

**Figure SI 2.**
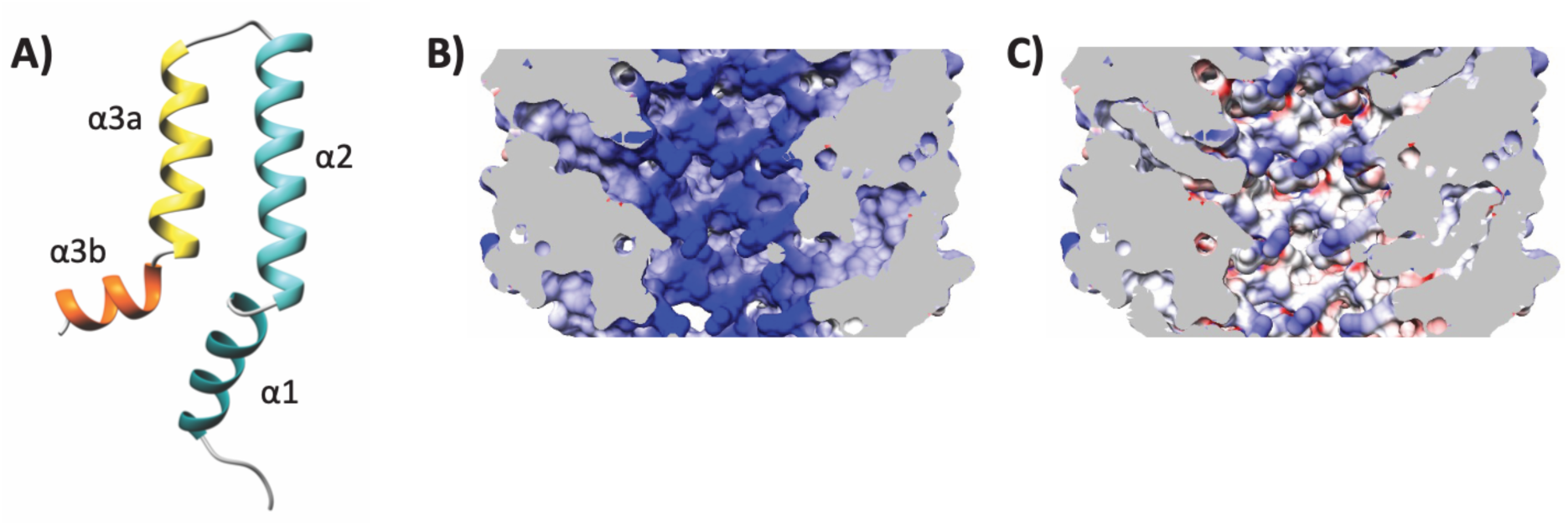
VirB2 monomer and Luminal electrostatic potential. (A) VirB2 monomer showing 4 helices (α1, α2, α3a and α3b). (B-C) Electrostatic potential of the T-pilus lumen shown (B) without lipid and (C) with lipid. Electrostatic potential of the lumen was calculated using Chimera.

**Figure SI 3.**
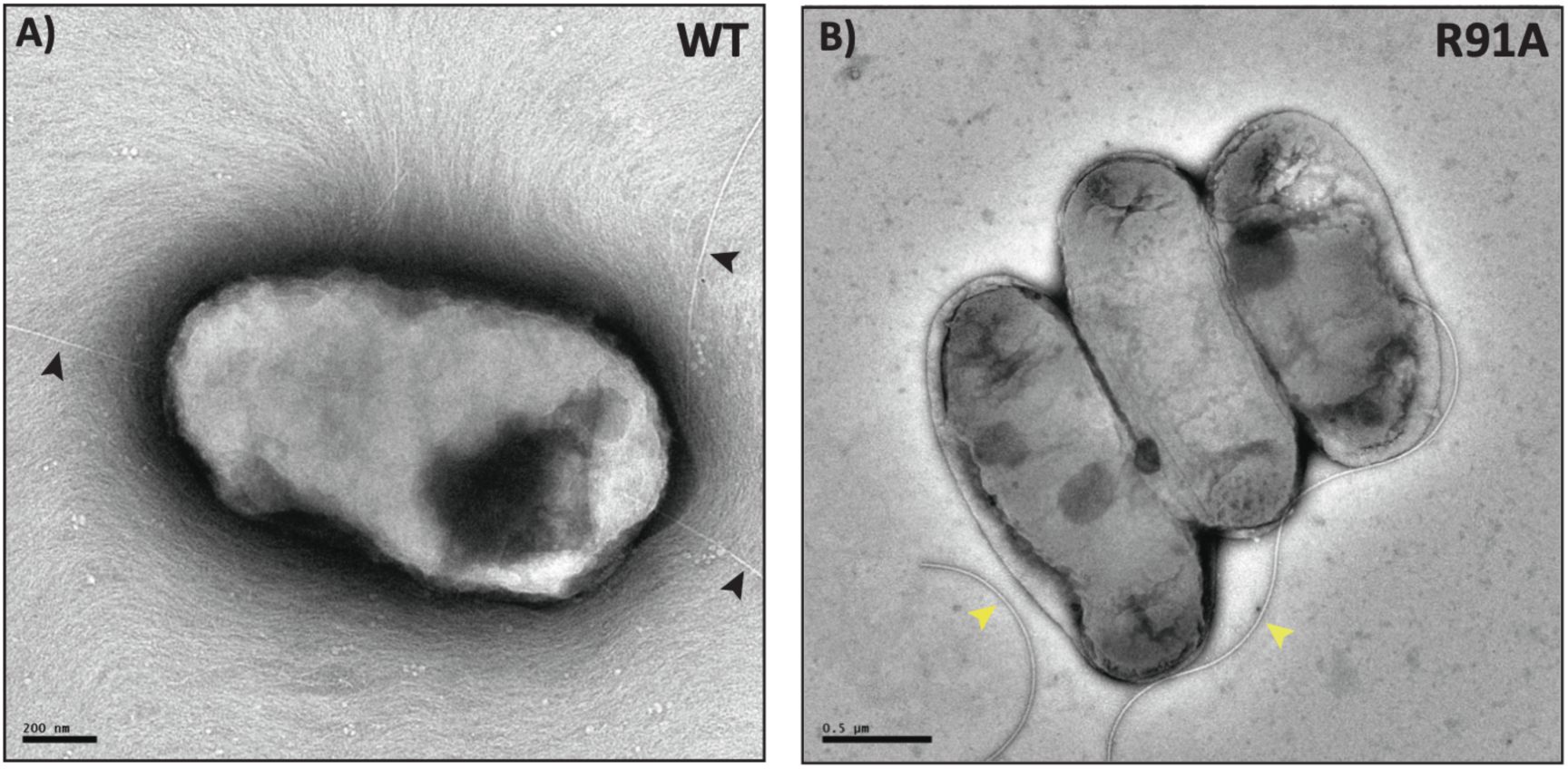
Effect of R91 mutation on T-pilus formation. Negatively stained micrographs of (A) WT and (B) R91A mutants showing presence of T-pili and absence of T-pilus, respectively. Black arrows show T-pili (∼10 nm in diameter), yellow arrows show flagella. Scale bar, as shown on the micrographs.

**Figure SI 4.**
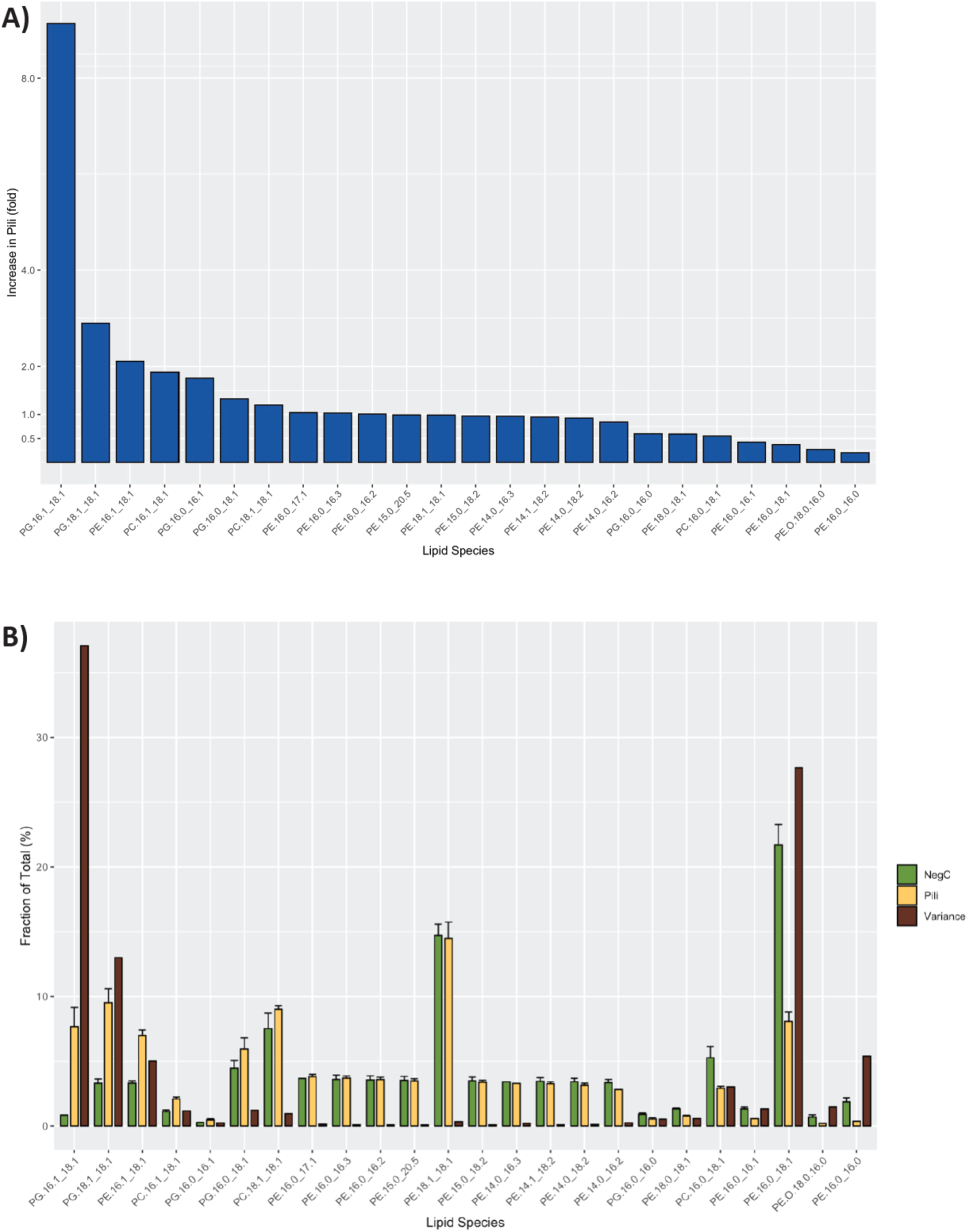
Lipidomic analysis the T-pilus. (A) Barplot showing the fold increase of each analyzed lipid species in the pili-containing sample compared to the negative control (B) Barplot showing relative abundances of lipid species (in percent) in purified pilus sample (yellow) and negative control (green), as an average of three independent experiments. The bars are ordered by the fold increase in pili. The variance was calculated for each lipid species for all six observations (three from pilus, and three from negative control) on CLR-transformed data. This value was then divided by the summed variance of all species. This Contribution to Total Variance (in percent) for each lipid species is shown in brown.

**Table SI 1.**
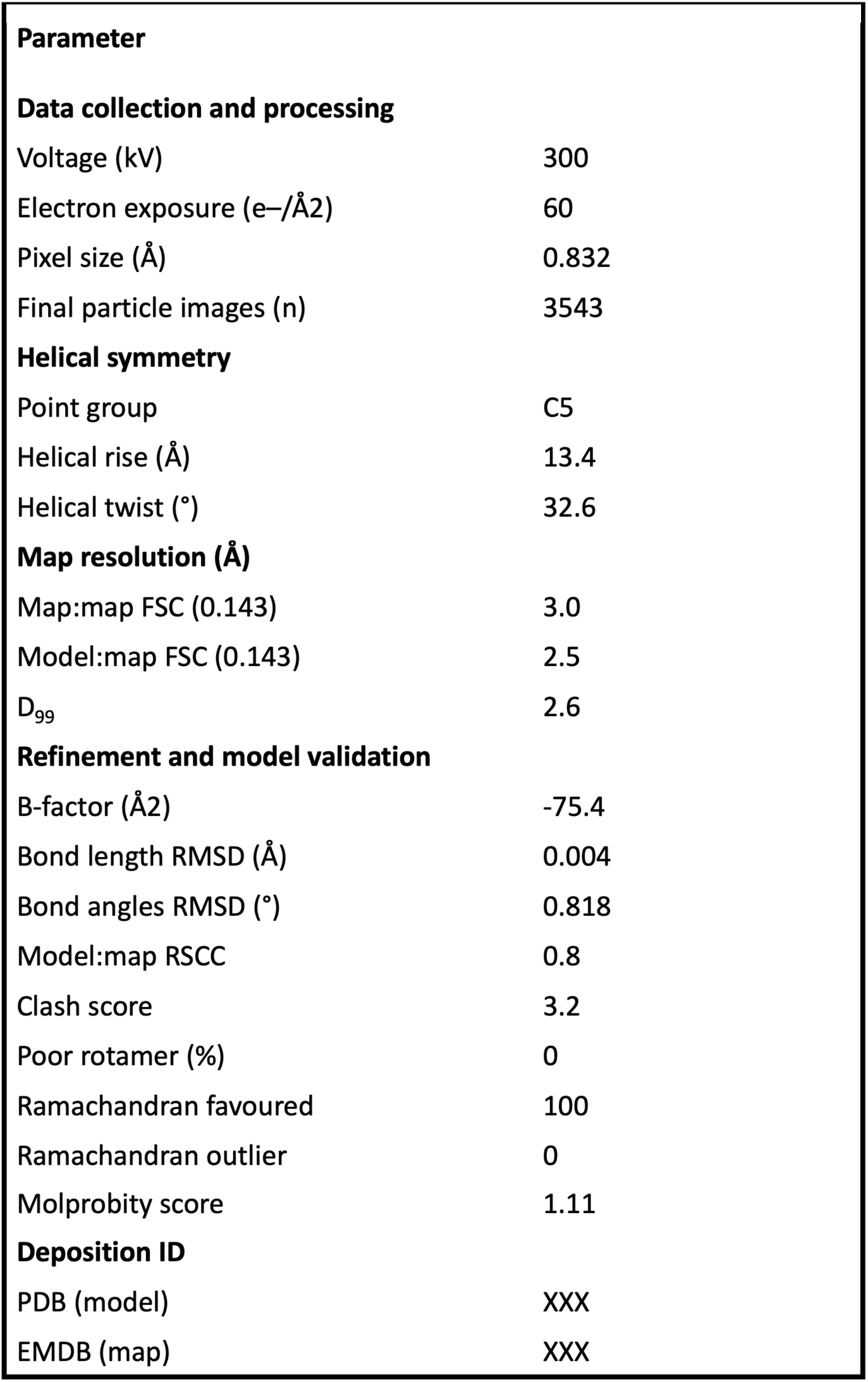
Cryo-EM data collection, refinement and validation statistics.

| Table 1. Cryo-EM data collection, refinement and validation statistics |  |
| --- | --- |
| Parameter |  |
| Data collection and processing |  |
| Voltage (kV) | 300 |
| Electron exposure (e<sup>-</sup>/Å<sup>2</sup>) | 60 |
| Pixel size (Å) | 0.832 |
| Final particle images (n) | 3543 |
| Helical symmetry |  |
| Point group | C5 |
| Helical rise (Å) | 13.4 |
| Helical twist (°) | 32.6 |
| Map resolution (Å) |  |
| Map:map FSC (0.143) | 3.0 |
| Model:map FSC (0.143) | 2.5 |
| D<sub>99</sub> | 2.6 |
| Refinement and model validation |  |
| B-factor (Å<sup>2</sup>) | -75.4 |
| Bond length RMSD (Å) | 0.004 |
| Bond angles RMSD (°) | 0.818 |
| Model:map RSCC | 0.8 |
| Clash score | 3.2 |
| Poor rotamer (%) | 0 |
| Ramachandran favoured | 100 |
| Ramachandran outlier | 0 |
| Molprobity score | 1.11 |
| Deposition ID |  |
| PDB (model) | XXX |
| EMDB (map) | XXX |

**Table SI 2.**
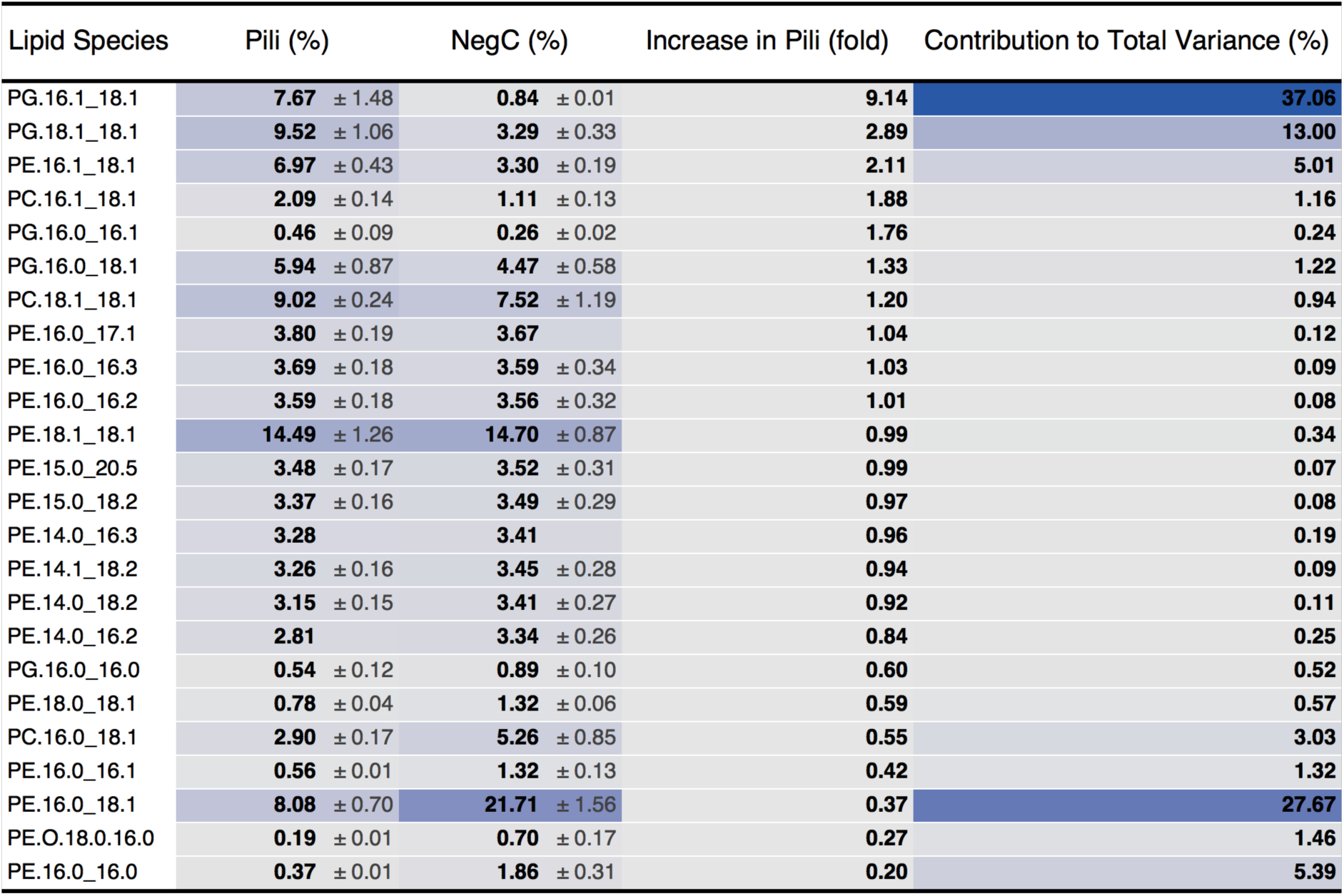
Lipidomic analysis of the T-pilus. Relative abundances of lipid species with SD in pili-containing samples and negative control as an average of three independent preparations.

| Lipid Species | Pili (%) | NegC (%) | Increase in Pili (fold) | Contribution to Total Variance (%) |
| --- | --- | --- | --- | --- |
| PG.16.1_18.1 | **7.67** ± 1.48 | **0.84** ± 0.01 | **9.14** | **37.06** |
| PG.18.1_18.1 | **9.52** ± 1.06 | **3.29** ± 0.33 | **2.89** | **13.00** |
| PE.16.1_18.1 | **6.97** ± 0.43 | **3.30** ± 0.19 | **2.11** | **5.01** |
| PC.16.1_18.1 | **2.09** ± 0.14 | **1.11** ± 0.13 | **1.88** | **1.16** |
| PG.16.0_16.1 | **0.46** ± 0.09 | **0.26** ± 0.02 | **1.76** | **0.24** |
| PG.16.0_18.1 | **5.94** ± 0.87 | **4.47** ± 0.58 | **1.33** | **1.22** |
| PC.18.1_18.1 | **9.02** ± 0.24 | **7.52** ± 1.19 | **1.20** | **0.94** |
| PE.16.0_17.1 | **3.80** ± 0.19 | **3.67** | **1.04** | **0.12** |
| PE.16.0_16.3 | **3.69** ± 0.18 | **3.59** ± 0.34 | **1.03** | **0.09** |
| PE.16.0_16.2 | **3.59** ± 0.18 | **3.56** ± 0.32 | **1.01** | **0.08** |
| PE.18.1_18.1 | **14.49** ± 1.26 | **14.70** ± 0.87 | **0.99** | **0.34** |
| PE.15.0_20.5 | **3.48** ± 0.17 | **3.52** ± 0.31 | **0.99** | **0.07** |
| PE.15.0_18.2 | **3.37** ± 0.16 | **3.49** ± 0.29 | **0.97** | **0.08** |
| PE.14.0_16.3 | **3.28** | **3.41** | **0.96** | **0.19** |
| PE.14.1_18.2 | **3.26** ± 0.16 | **3.45** ± 0.28 | **0.94** | **0.09** |
| PE.14.0_18.2 | **3.15** ± 0.15 | **3.41** ± 0.27 | **0.92** | **0.11** |
| PE.14.0_16.2 | **2.81** | **3.34** ± 0.26 | **0.84** | **0.25** |
| PG.16.0_16.0 | **0.54** ± 0.12 | **0.89** ± 0.10 | **0.60** | **0.52** |
| PE.18.0_18.1 | **0.78** ± 0.04 | **1.32** ± 0.06 | **0.59** | **0.57** |
| PC.16.0_18.1 | **2.90** ± 0.17 | **5.26** ± 0.85 | **0.55** | **3.03** |
| PE.16.0_16.1 | **0.56** ± 0.01 | **1.32** ± 0.13 | **0.42** | **1.32** |
| PE.16.0_18.1 | **8.08** ± 0.70 | **21.71** ± 1.56 | **0.37** | **27.67** |
| PE.O.18.0.16.0 | **0.19** ± 0.01 | **0.70** ± 0.17 | **0.27** | **1.46** |
| PE.16.0_16.0 | **0.37** ± 0.01 | **1.86** ± 0.31 | **0.20** | **5.39** |

## Notes

### Competing Interest Statement

The authors have declared no competing interest.

## References

Adams, P.D., Afonine, P. V, Bunkóczi, G., Chen, V.B., Echols, N., Headd, J.J., Hung, L.-W., Jain, S., Kapral, G.J., and Kunstleve, R.W.G. (2011). The Phenix software for automated determination of macromolecular structures. Methods 55, 94–106..

Aktas, M., Danne, L., Möller, P., and Narberhaus, F. (2014). Membrane lipids in Agrobacterium tumefaciens: Biosynthetic pathways and importance for pathogenesis. Front. Plant Sci. 5, 1–13. https://doi.org/10.3389/fpls.2014.00109.

Aly, K.A., and Baron, C. (2007). The VirB5 protein localizes to the T-pilus tips in Agrobacterium tumefaciens. Microbiology 153, 3766–3775. https://doi.org/10.1099/mic.0.2007/010462-0.

Amro, J., Black, C., Jemouai, Z., Rooney, N., Daneault, C., Zeytuni, N., Ruiz, M., Bui, K.H., and Baron, C. (2022). Cryo-EM structure of the *Agrobacterium tumefaciens* T-pilus reveals the importance of positive charges in the lumen. BioRxiv 2022.04.28.489814. https://doi.org/10.1101/2022.04.28.489814.

Van Attikum, H., Bundock, P., and Hooykaas, P.J.J. (2001). Non-homologous end-joining proteins are required for Agrobacterium T-DNA integration. EMBO J. 20, 6550–6558. https://doi.org/10.1093/emboj/20.22.6550.

Beijersbergen, A., Smith, S.J., and Hooykaas, P.J.J. (1994). Localization and Topology of VirB Proteins of Agrobacterium tumefaciens. Plasmid 32, 212–218. https://doi.org/https://doi.org/10.1006/plas.1994.1057.

Berger, B.R., and Christie, P.J. (1994). Genetic complementation analysis of the Agrobacterium tumefaciens virB operon: virB2 through virB11 are essential virulence genes. J. Bacteriol. 176, 3646– 3660. .

Chesnokova, O., Coutinho, J.B., Khan, I.H., Mikhail, M.S., and Kado, C.I. (1997). Characterization of flagella genes of Agrobacterium tumefaciens, and the effect of a bald strain on virulence. Mol. Microbiol. 23, 579–590. https://doi.org/10.1046/j.1365-2958.1997.d01-1875.x.

Christie, P.J. (2016). The Mosaic Type IV Secretion Systems. EcoSal Plus 7. https://doi.org/10.1128/ecosalplus.ESP-0020-2015.

Costa, T.R.D., Ilangovan, A., Ukleja, M., Redzej, A., Santini, J.M., Smith, T.K., Egelman, E.H., and Waksman, G. (2016). Structure of the Bacterial Sex F Pilus Reveals an Assembly of a Stoichiometric Protein-Phospholipid Complex. Cell 166, 1436–1444.e10. https://doi.org/10.1016/j.cell.2016.08.025.

Costa, T.R.D., Harb, L., Khara, P., Zeng, L., Hu, B., and Christie, P.J. (2021). Type IV secretion systems: advances in structure, function, and activation. Mol. Microbiol. 115, 436–452..

Croll, T.I. (2018). ISOLDE: a physically realistic environment for model building into low-resolution electron-density maps. Acta Crystallogr. Sect. D Struct. Biol. 74, 519–530..

Dessaux, Y., Petit, A., Farrand, S.K., and Murphy, P.J. (1998). Opines and Opine-Like Molecules Involved in Plant-Rhizobiaceae Interactions BT - The Rhizobiaceae: Molecular Biology of Model Plant-Associated Bacteria. H.P. Spaink, A. Kondorosi, and P.J.J. Hooykaas, eds. (Dordrecht: Springer Netherlands), pp. 173–197.

Eisenbrandt, R., Kalkum, M., Lai, E.M., Lurz, R., Kado, C.I., and Lanka, E. (1999). Conjugative pili of IncP plasmids, and the Ti plasmid T pilus are composed of cyclic subunits. J. Biol. Chem. 274, 22548– 22555. https://doi.org/10.1074/jbc.274.32.22548.

Emsley, P., Lohkamp, B., Scott, W.G., and Cowtan, K. (2010). Features and development of Coot. Acta Crystallogr. Sect. D Biol. Crystallogr. 66, 486–501..

Erh-Min, L., and I., K.C. (1998). Processed VirB2 Is the Major Subunit of the Promiscuous Pilus of Agrobacterium tumefaciens. J. Bacteriol. 180, 2711–2717. https://doi.org/10.1128/JB.180.10.2711-2717.1998.

Frost, L.S., Armstrong, G.D., Finlay, B.B., Edwards, B.F., and Paranchych, W. (1983). N-terminal amino acid sequencing of EDP208 conjugative pili. J. Bacteriol. 153, 950–954..

Frost, L.S., Paranchych, W., and Willetts, N.S. (1984). DNA sequence of the F traALE region that includes the gene for F pilin. J. Bacteriol. 160, 395–401. https://doi.org/10.1128/jb.160.1.395-401.1984.

Goddard, T.D., Huang, C.C., Meng, E.C., Pettersen, E.F., Couch, G.S., Morris, J.H., and Ferrin, T.E. (2018). UCSF ChimeraX: Meeting modern challenges in visualization and analysis. Protein Sci. 27, 14–25..

Gordon, J.E., Costa, T.R.D., Patel, R.S., Gonzalez-Rivera, C., Sarkar, M.K., Orlova, E. V., Waksman, G., and Christie, P.J. (2017). Use of chimeric type IV secretion systems to define contributions of outer membrane subassemblies for contact-dependent translocation. Mol. Microbiol. 105, 273–293. https://doi.org/10.1111/mmi.13700.

Greenacre, M. (2021). Compositional data analysis. Annu. Rev. Stat. Its Appl. 8, 271–299..

Groenewold, M.K., Hebecker, S., Fritz, C., Czolkoss, S., Wiesselmann, M., Heinz, D.W., Jahn, D., Narberhaus, F., Aktas, M., and Moser, J. (2019). Virulence of Agrobacterium tumefaciens requires lipid homeostasis mediated by the lysyl-phosphatidylglycerol hydrolase AcvB. Mol. Microbiol. 111, 269–286..

Guex, N., Peitsch, M.C., and Schwede, T. (2009). Automated comparative protein structure modeling with SWISS-MODEL and Swiss-PdbViewer: A historical perspective. Electrophoresis 30, S162–S173..

He, S., and Scheres, S.H.W. (2017). Helical reconstruction in RELION. J. Struct. Biol. 198, 163–176. https://doi.org/10.1016/j.jsb.2017.02.003.

Hsieh, W.-Y., Williams, K.J., Su, B., and Bensinger, S.J. (2021). Profiling of mouse macrophage lipidome using direct infusion shotgun mass spectrometry. STAR Protoc. 2, 100235..

Hwang, H.-H., Yu, M., and Lai, E.-M. (2017). *Agrobacterium*-Mediated Plant Transformation: Biology and Applications. Arab. B. 2017. https://doi.org/10.1199/tab.0186.

Idnurm, A., Bailey, A.M., Cairns, T.C., Elliott, C.E., Foster, G.D., Ianiri, G., and Jeon, J. (2017). A silver bullet in a golden age of functional genomics: The impact of agrobacterium-mediated transformation of fungi. Fungal Biol. Biotechnol. 4, 1–28. https://doi.org/10.1186/s40694-017-0035-0.

Jakubowski, S.J., Krishnamoorthy, V., Cascales, E., and Christie, P.J. (2004). Agrobacterium tumefaciens VirB6 domains direct the ordered export of a DNA substrate through a type IV secretion system. J. Mol. Biol. 341, 961–977. https://doi.org/10.1016/j.jmb.2004.06.052.

Jakubowski, S.J., Cascales, E., Krishnamoorthy, V., and Christie, P.J. (2005). Agrobacterium tumefaciens VirB9, an outer-membrane-associated component of a type IV secretion system, regulates substrate selection and T-pilus biogenesis. J. Bacteriol. 187, 3486–3495. https://doi.org/10.1128/JB.187.10.3486-3495.2005.

Jakubowski, S.J., Kerr, J.E., Garza, I., Krishnamoorthy, V., Bayliss, R., Waksman, G., and Christie, P.J. (2009). Agrobacterium VirB10 domain requirements for type IV secretion and T pilus biogenesis. Mol. Microbiol. 71, 779–794. https://doi.org/https://doi.org/10.1111/j.1365-2958.2008.06565.x.

Kalogeraki, V.S., and Winans, S.C. (1995). The octopine-type Ti plasmid pTiA6 of Agrobacterium tumefaciens contains a gene homologous to the chromosomal virulence gene acvB. J. Bacteriol. 177, 892–897. https://doi.org/10.1128/jb.177.4.892-897.1995.

Kerr, J.E., and Christie, P.J. (2010). Evidence for VirB4-mediated dislocation of membrane-integrated VirB2 pilin during biogenesis of the Agrobacterium VirB/VirD4 type IV secretion system. J. Bacteriol. 192, 4923–4934..

Klüsener, S., Hacker, S., Tsai, Y.L., Bandow, J.E., Gust, R., Lai, E.M., and Narberhaus, F. (2010). Proteomic and transcriptomic characterization of a virulence-deficient phosphatidylcholine-negative Agrobacterium tumefaciens mutant. Mol. Genet. Genomics 283, 575–589. https://doi.org/10.1007/s00438-010-0542-7.

Kuldau, G.A., De Vos, G., Owen, J., McCaffrey, G., and Zambryski, P. (1990). The virB operon of Agrobacterium tumefaciens pTiC58 encodes 11 open reading frames. Mol. Gen. Genet. MGG 221, 256–266. https://doi.org/10.1007/BF00261729.

Lai, E., and Kado, C.I. (2000). The T-pilus of Agrobacterium tumefaciens. 361–369..

Lai, E.M., and Kado, C.I. (2002). The Agrobacterium tumefaciens T pilus composed of cyclic T pilin is highly resilient to extreme environments. FEMS Microbiol. Lett. 210, 111–114. https://doi.org/10.1016/S0378-1097(02)00601-8.

Leaver-Fay, A., Tyka, M., Lewis, S.M., Lange, O.F., Thompson, J., Jacak, R., Kaufman, K.W., Renfrew, P.D., Smith, C.A., and Sheffler, W. (2011). ROSETTA3: an object-oriented software suite for the simulation and design of macromolecules. In Methods in Enzymology, (Elsevier), pp. 545–574.

Lederberg, J., and Tatum, E.L. (1946). Gene recombination in Escherichia coli. Nature 158, 558..

Li, Y.G., and Christie, P.J. (2018). The Agrobacterium VirB/VirD4 T4SS: mechanism and architecture defined through in vivo mutagenesis and chimeric systems. Agrobacterium Biol. 233–260..

Macé, K., Vadakkepat, A.K., Redzej, A., Lukoyanova, N., Oomen, C., Braun, N., Ukleja, M., Lu, F., Costa, T.R.D., Orlova, E. V., et al. (2022). Cryo-EM structure of a type IV secretion system. Nature 607. https://doi.org/10.1038/s41586-022-04859-y.

María, G., and Fernando, de la C. (2018). Natural and Artificial Strategies To Control the Conjugative Transmission of Plasmids. Microbiol. Spectr. 6, 6.1.03. https://doi.org/10.1128/microbiolspec.MTBP-0015-2016.

McCullen, C.A., and Binns, A.N. (2006). Agrobacterium tumefaciens and plant cell interactions and activities required for interkingdom macromolecular transfer. Annu. Rev. Cell Dev. Biol. 22, 101–127. https://doi.org/10.1146/annurev.cellbio.22.011105.102022.

Meng, R., Jiang, M., Cui, Z., Chang, J.Y., Yang, K., Jakana, J., Yu, X., Wang, Z., Hu, B., and Zhang, J. (2019). Structural basis for the adsorption of a single-stranded RNA bacteriophage. Nat. Commun. 10. https://doi.org/10.1038/s41467-019-11126-8.

Mossey, P., Hudacek, A., and Das, A. (2010). Agrobacterium tumefaciens type IV secretion protein VirB3 is an inner membrane protein and requires VirB4, VirB7, and VirB8 for stabilization. J. Bacteriol. 192, 2830–2838..

Norman, A., Hansen, L.H., and Sørensen, S.J. (2009). Conjugative plasmids: vessels of the communal gene pool. Philos. Trans. R. Soc. B Biol. Sci. 364, 2275–2289. https://doi.org/10.1098/rstb.2009.0037.

Palarea-Albaladejo, J., and Martín-Fernández, J.A. (2015). zCompositions — R package for multivariate imputation of left-censored data under a compositional approach. Chemom. Intell. Lab. Syst. 143, 85–96. https://doi.org/https://doi.org/10.1016/j.chemolab.2015.02.019.

Pan, S.Q., Jin, S., Boulton, M.I., Hawes, M., Gordon, M.P., and Nester, E.W. (1995). An Agrobacterium virulence factor encoded by a Ti plasmid gene or a chromosomal gene is required for T-DNA transfer into plants. Mol. Microbiol. 17, 259–269. https://doi.org/https://doi.org/10.1111/j.1365-2958.1995.mmi_17020259.x.

Phillips, J.C., Hardy, D.J., Maia, J.D.C., Stone, J.E., Ribeiro, J. V, Bernardi, R.C., Buch, R., Fiorin, G., Hénin, J., and Jiang, W. (2020). Scalable molecular dynamics on CPU and GPU architectures with NAMD. J. Chem. Phys. 153, 44130. .

R Core Team, R.C. (2013). R: A language and environment for statistical computing. R Foundation for Statistical Computing, Vienna, Austria. Http//Www.R-Project.Org/.

Rohou, A., and Grigorieff, N. (2015). CTFFIND4: Fast and accurate defocus estimation from electron micrographs. J. Struct. Biol. 192, 216–221. https://doi.org/10.1016/j.jsb.2015.08.008.

Scheres, S.H.W. (2012). RELION: Implementation of a Bayesian approach to cryo-EM structure determination. J. Struct. Biol. 180, 519–530. https://doi.org/https://doi.org/10.1016/j.jsb.2012.09.006.

Schmidt-Eisenlohr, H., Domke, N., Angerer, C., Wanner, G., Zambryski, P.C., and Baron, C. (1999). Vir proteins stabilize VirB5 and mediate its association with the T pilus of Agrobacterium tumefaciens. J. Bacteriol. 181, 7485–7492. https://doi.org/10.1128/jb.181.24.7485-7492.1999.

Shirasu, K., and Kado, C.I. (1993). Membrane location of the Ti plasmid VirB proteins involved in the biosynthesis of a pilin-like conjugative structure on Agrobacterium tumefaciens. FEMS Microbiol. Lett. 111, 287–293. https://doi.org/10.1111/j.1574-6968.1993.tb06400.x.

Shirasu, K., Morel, P., and Kado, C.I. (1990). Characterization of the virB operon of an Agrobacterium tumefaciens Ti plasmid: nucleotide sequence and protein analysis. Mol. Microbiol. 4, 1153–1163. https://doi.org/https://doi.org/10.1111/j.1365-2958.1990.tb00690.x.

Su, B., Bettcher, L.F., Hsieh, W.-Y., Hornburg, D., Pearson, M.J., Blomberg, N., Giera, M., Snyder, M.P., Raftery, D., and Bensinger, S.J. (2021). A DMS shotgun lipidomics workflow application to facilitate high-throughput, comprehensive lipidomics. J. Am. Soc. Mass Spectrom. 32, 2655–2663..

Vinuesa, P., Neumann-Silkow, F., Pacios-Bras, C., Spaink, H.P., Martínez-Romero, E., and Werner, D. (2003). Genetic analysis of a pH-regulated operon from Rhizobium tropici CIAT899 involved in acid tolerance and nodulation competitiveness. Mol. Plant-Microbe Interact. 16, 159–168. https://doi.org/10.1094/MPMI.2003.16.2.159.

Virolle, C., Goldlust, K., Djermoun, S., Bigot, S., and Lesterlin, C. (2020). Plasmid transfer by conjugation in gram-negative bacteria: From the cellular to the community level. Genes (Basel). 11, 1–33. https://doi.org/10.3390/genes11111239.

Wickham, H. (2016). Data analysis. In Ggplot2, (Springer), pp. 189–201.

Winans, S.C. (1990). Transcriptional induction of an Agrobacterium regulatory gene at tandem promoters by plant-released phenolic compounds, phosphate starvation, and acidic growth media. J. Bacteriol. 172, 2433–2438..

Wirawan, I.G.P., Hi Wan Kang, and Kojima, M. (1993). Isolation and characterization of a new chromosomal virulence gene of Agrobacterium tumefaciens. J. Bacteriol. 175, 3208–3212. https://doi.org/10.1128/jb.175.10.3208-3212.1993.

Wu, H.Y., Chen, C.Y., and Lai, E.M. (2014). Expression and functional characterization of the Agrobacterium VirB2 amino acid substitution variants in T-pilus biogenesis, virulence, and transient transformation efficiency. PLoS One 9, 1–11. https://doi.org/10.1371/journal.pone.0101142.

Zheng, S.Q., Palovcak, E., Armache, J.P., Verba, K.A., Cheng, Y., and Agard, D.A. (2017). MotionCor2: Anisotropic correction of beam-induced motion for improved cryo-electron microscopy. Nat. Methods 14, 331–332. https://doi.org/10.1038/nmeth.4193.

Zheng, W., Pena, A., Low, W.W., Wong, J.L.C., Frankel, G., and Egelman, E.H. (2020). Cryoelectron-Microscopic Structure of the pKpQIL Conjugative Pili from Carbapenem-Resistant Klebsiella pneumoniae. Structure 28, 1321–1328.e2. https://doi.org/https://doi.org/10.1016/j.str.2020.08.010.

Zhu, J., Oger, P.M., Schrammeijer, B., Hooykaas, P.J.J., Farrand, S.K., and Winans, S.C. (2000). The bases of crown gall tumorigenesis. J. Bacteriol. 182, 3885–3895..

